# Using summary statistics to evaluate the genetic architecture of multiplicative combinations of initially analyzed phenotypes with a flexible choice of covariates

**DOI:** 10.1101/2021.03.08.433979

**Authors:** Jack Wolf, Jason Westra, Nathan Tintle

## Abstract

While the promise of electronic medical record and biobank data is large, major questions remain about patient privacy, computational hurdles, and data access. One promising area of recent development is pre-computing non-individually identifiable summary statistics to be made publicly available for exploration and downstream analysis. In this manuscript we demonstrate how to utilize pre-computed linear association statistics between individual genetic variants and phenotypes to infer genetic relationships between products of phenotypes (e.g., ratios; logical combinations of binary phenotypes using ‘and’ and ‘or’) with customized covariate choices. We propose a method to approximate covariate adjusted linear models for products and logical combinations of phenotypes using only pre-computed summary statistics. We evaluate our method’s accuracy through several simulation studies and an application modeling various fatty acid ratios using data from the Framingham Heart Study. These studies show consistent ability to recapitulate analysis results performed on individual level data including maintenance of the Type I error rate, power, and effect size estimates. An implementation of this proposed method is available in the publicly available R package pcsstools.

## 1 Introduction

Researchers now have readily available access to massive quantities of genotypic and phenotypic data (Cox, 2018; Simell et al., 2019). For example, via the Electronic Medical Records and Genomics (eMERGE Network; https://www.genome.gov/Funded-Programs-Projects/Electronic-Medical-Records-and-Genomics-Network-eMERGE, the UK-Biobank (Bycroft et al., 2018) other initiatives and repositories (e.g., 23andMe, MGI http://pheweb.sph.umich.edu/ (Gagliano Taliun et al., 2020), FINRISK, CHOP (Diogo et al., 2018), among others), researchers can access a wide variety of phenotypic and genomics data on hundreds of thousands of individuals. However, important questions remain about how to best leverage these repositories. For example, the size of biobank datasets makes it challenging to transfer, store, and analyze data locally. While cloud computing minimizes some of these issues, it brings its own challenges related to cost (storage and computation), transfer, and access. Furthermore, data security and privacy issues are of paramount importance throughout all aspects of the data access, storage, and analysis pipeline (Heatherly, 2016; Jones et al., 2012; Simell et al., 2019).

A key innovation in this field is precomputing non-individually identifiable summary statistics on biobank data and maximizing access to this data (Pasaniuc & Price, 2017). For example, GeneAtlas provides basic summary statistics for simple linear regression models of single nucleotide variants (SNVs) with 1000s of available phenotypic variables across hundreds of thousands of individuals in the UK Biobank (Canela-Xandri et al., 2018), which also provides access to phenotype-phenotype correlations, single nucleotide polymorphism (SNP) minor allele frequencies (MAFs) and Hardy Weinberg Equilibrium (HWE) *p*-values. Likewise, PheWeb is a software toolkit which provides access to the UK Biobank and Michigan Genomics Initiative data via a series of easy-to-navigate visualization and summary tools (http://pheweb.sph.umich.edu/)(Gagliano Taliun et al., 2020; Neale, B. M., 2018). Others simply provide access to sets of pre-computed summary statistics (PCSS) from large datasets (e.g., https://www.leelabsg.org/resources). These resources mitigate many of the privacy and security concerns mentioned above since no individual participant data (IPD) is shared. In addition, the size of these repositories are only fractions of the size of IPD, making transfer and storage of the data much more efficient. Finally, these services provide PCSS, which alleviates much of the computational burden on researchers. Despite these advantages, significant limitations currently exist when using these repositories of PCSS.

For example, researchers may want to modify a phenotype with available PCSS to one that is of greater clinical interest or use different sets of covariates than those considered in pre-computed analyses. Recent work is beginning to address these limitations. In two recent papers by our group (Gasdaska et al., 2019; Wolf et al., 2020), we demonstrated how to use standard PCSS (only means, variances, and correlations of all predictors and responses) to calculate the coefficients and standard errors for the linear model for a linear combination of phenotypes with an arbitrary set of covariates. This can then be used to perform Principal Component Analysis (PCA) on a set of phenotypes since principal component scores are just linear combinations with weights derived from the phenotype covariance matrix. Further, we demonstrated that if the phenotype correlation matrix is not available, we can use the correlation of test statistics for each phenotype across all genetic markers in its place with little loss of efficiency. These innovations mean that researchers can, using only PCSS, select the unique set of covariates they wish to adjust for and model a linear combination of phenotypes.

Importantly, these two approaches which require a priori specification of a phenotype of clinical interest, contrast to other recently developed methods which jointly and simultaneously analyze multiple phenotypes (Dutta, Gagliano Taliun, et al., 2019; Dutta, Scott, et al., 2019; Guo & Wu, 2019; Li et al., 2020; Ray & Boehnke, 2018) without an explicit characterization of the relationship between the phenotypes. These joint phenotype tests aim to simultaneously analyze multiple phenotypes while satisfying statistical objectives such as maximizing power under certain conditions. Furthermore, some of these approaches (Guo & Wu, 2019; Ray & Boehnke, 2018) do so using PCSS readily available from existing repositories.

Currently, our group’s methods for using PCSS to analyze modified phenotypes with flexible covariate choices are limited to PCA and choosing a phenotype that is a linear combination of the phenotypes for which PCSS are available. In this manuscript, we demonstrate how to analyze modified phenotypes which are multiplicative combinations of an arbitrarily large number of phenotypes for which PCSS are available. We also demonstrate how to flexibly adjust for covariates in these modified phenotype models. Importantly, we also show how the multiplication of phenotypes, when applied to binary phenotypes, allows for logical combination (“and” and “or”) of phenotypes (e.g., to do inference on a phenotype ***y***, that is “***y***_1_ or ***y***_2_”). After presenting a mathematical framework for the method, we validate the method using comprehensive simulations and demonstrate the method on real data from the Framingham Heart Study.

## 2 Methods

Consider the *m* phenotypes ***y***_1_, …, ***y***_*m*_ where each is an *n*×1 vector of measures across *n* subjects and the *n*×*p* design matrix ***X*** = (***x***_1_, …, ***x***_*p*_) which consists of variables including genotypic information, covariates, and an intercept column. Moreover, let ***w***_*m*_ = ***y***_1_***y***_2_ · ***y***_*m*_ denote the pairwise Hadamard product of all *m* phenotypes for each subject. Our aim is to approximate the coefficients and standard errors of the covariate adjusted linear regression model for the product of *m* phenotypes: 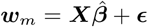 using only readily available PCSS.

### 2.1 Assumed Pre-Computed Summary Statistics and Information

As is typically made available, we assume knowledge of the following PCSS: the means of every predictor (e.g. SNPs and covariates), the means of every phenotype, and the full variance-covariance matrix of all predictors and phenotypes (i.e. 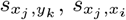 and 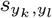 for any *i, j, k, l* where 1 ≤ *i, j* ≤ *p* and 1 ≤ *k, l* ≤ *m*). These are all readily available in standard PCSS repositories. We also assume to know the distribution that each predictor and phenotype follows (e.g. binary, log-normal, etc.). Figure 1 displays the assumed information when modeling via both IPD and PCSS.

**Figure 1:**
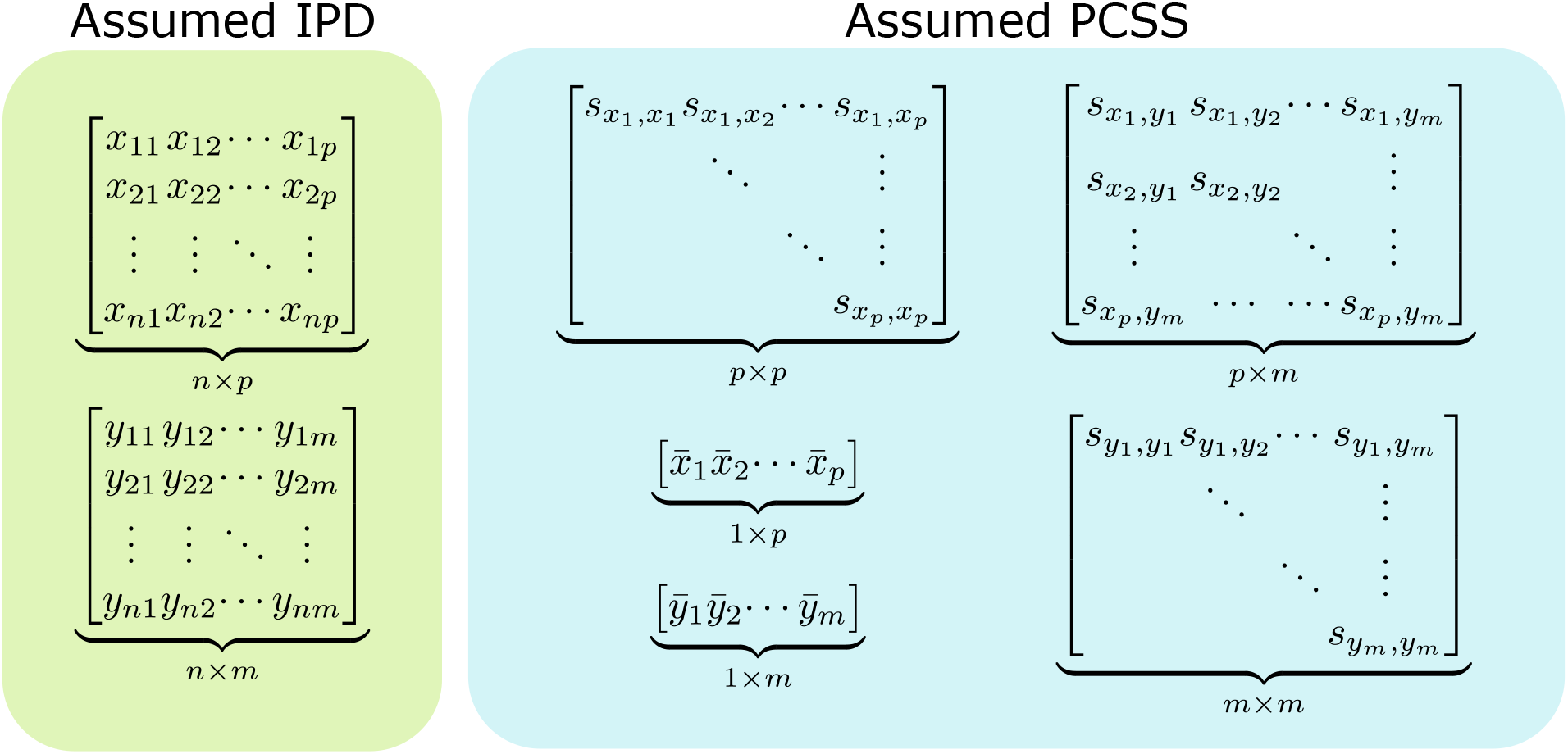
Data assumed when modeling using individual participant data (IPD) and when using pre-computed summary statistics (PCSS) to model a product of *m* phenotypes (***y***_1_, …, ***y***_*m*_) as a linear function of *p* covariates (***x***_1_, …, ***x***_*m*_). While modeling using IPD requires *n* × (*p* + *m*) points of data, using PCSS only requires *p*^2^ + *pm* + *m*^2^ + *p* + *m* values, which is far less when *n* is moderately large compared to *p* and *m*. All of these PCSS are readily available in existing PCSS repositories, or can be derived or approximated from other PCSS.

However, if some summary statistics are unknown, they may be able to be derived or approximated. For example, SNPs distributed in HWE can have their mean and variance approximated through a binomial distribution given the MAF. Furthermore, the covariance of a genetic variant and a non-genetic variable is calculated as the single-marker slope coefficient (for the model with the non-genetic variable as the response and the genetic variant as the predictor) divided by the variance of the genetic variant. Other published papers (Kim et al., 2015; Zhu et al., 2015) have shown that the correlation of two traits can be approximated by the correlation of *Z* statistics of SNPs not associated with either trait; i.e., 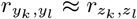 where ***z***_*k*_ and ***z***_*l*_ are vectors of single-marker test statistics for traits ***y***_*k*_ and ***y***_*l*_ across a genome wide association study filtered such that the associated *p*-values are above a set threshold for both traits. This approximation method is described in detail in Ray & Boehnke (2018). Two of our previous papers (Gasdaska et al., 2019; Wolf et al., 2020) have demonstrated the accuracy of these three methods through both simulation and real-data applications.

### 2.2 Linear Regression with Covariates using Pre-Computed Summary Statistics

Given a response vector ***w***_*m*_ and design matrix ***X*** = (***x***_1_, …, ***x***_*p*_) which includes *p* variables including SNPs’ minor allele counts, covariates, and a possible intercept column, the normal error regression model ***w***_*m*_ = ***Xβ*** + ***ϵ*** where ***ϵ*** ∼ *N* (**0**, *σ*^*2*^ ***I***) has ordinary least squares estimate for 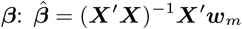. Further, Var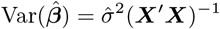. In a recent paper (Wolf et al., 2020), we demonstrated how to calculate these values using only PCSS:

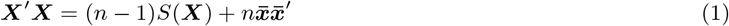

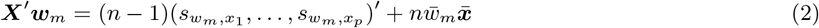

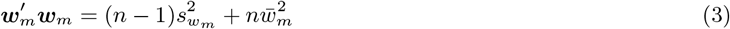

and

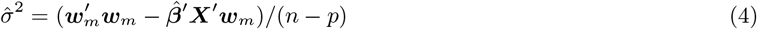

where *S*(***X***) is the *p* × *p* variance-covariance matrix of the columns of the design matrix ***X***, 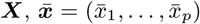 is the *p* × 1 vector of column means of ***X***, 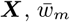 is the mean of ***w***_*m*_, and 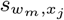 is the sample covariance between ***w***_*m*_ and ***x***_*j*_.

With these methods in mind and assumed access to standard PCSS, in order to approximate 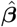, and 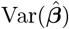 for this covariate adjusted multiple linear regression model, all that remains is to estimate 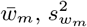 and 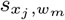 for each *j*. We will first demonstrate how to approximate these values when *m* = 2 and later show how recursion can be used to approximate covariances with *m* > 2 in Section 2.3.2.

### 2.3 Covariance Estimation

#### 2.3.1 Covariance Estimation with the Product of 2 Phenotypes

Let ***w***_2_ = ***y***_1_***y***_2_ be the pairwise Hadamard product of ***y***_1_ and ***y***_2_. Then, if ***x***_*j*_ represents an “intercept” column of the design matrix with all elements unity (i.e. if *x*_*j*_ = (1, …, 1)′), we set 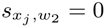. Otherwise, we proceed as follows:

We first approximate the conditional means and variances of ***y***_1_ and ***y***_2_ given ***x***_*j*_ = *x* through a linear regression model:

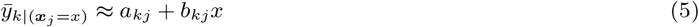

and

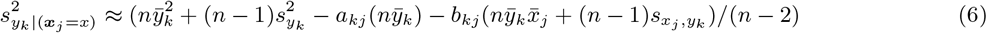

where 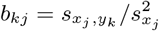 and 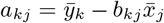. We note that this conditional variance will be constant at any value of ***x***_*j*_ following from the linear regression assumption of homoscedasticity.

Then, we calculate the sample partial correlation of ***y***_1_ and ***y***_2_ controlling for ***x***_*j*_ :

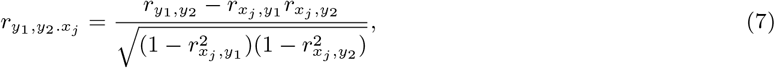

setting 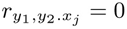 if either 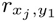 or 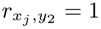. As the expectation of the conditional correlation equals the partial correlation under the assumption of a multivariate linear relationship between (***y***_1_, ***y***_2_) and ***x***_*j*_ (Baba et al., 2004), we use the partial correlation as an estimate of the conditional correlation of ***y***_1_ and ***y***_2_ at all possible values of ***x***_*j*_. So, we approximate the covariance of ***y***_1_ and ***y***_2_ conditional on ***x***_*j*_ :

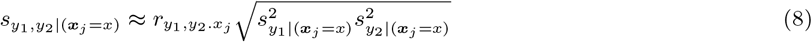

These terms let us approximate the conditional mean of ***w***_2_ at a given value *x* of ***x***_*j*_ :

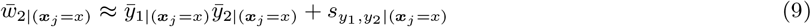

Then, letting *f*_*j*_ (*x*) be an assumed probability distribution /mass function for ***x***_*j*_ with support 𝒮_*j*_ (e.g. if ***x***_*j*_ is a vector of minor allele counts with MAF *p*, letting 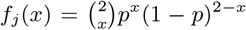 and 𝒮_*j*_ ={0,1,2}) we approximate the sample covariance of ***x***_*j*_ and ***w***_2_:

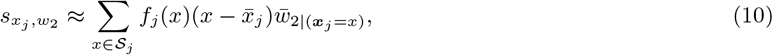

swapping the sums for integrals across the support when appropriate.

We calculate the sample mean of ***w***_2_ as

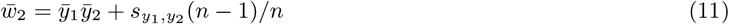

To approximate the variance, we first approximate the conditional variances of ***w***_2_ at all levels of ***x***_*j*_ :

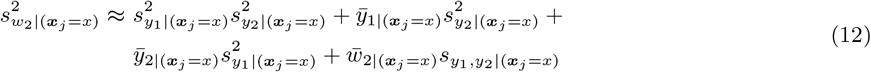

And then approximate the sample variance as:

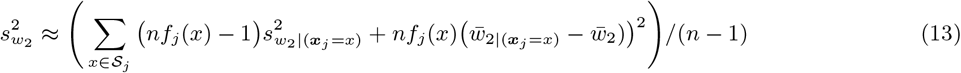

once again swapping the sum for an integral across 𝒮_*j*_ when appropriate. This approach leads to a different variance estimate for each predictor ***x***_*j*_. We treat the median of these estimates across each *j* as the estimated variance.

Hence, taking the means, variances, and pairwise covariances of ***x***_*j*_, ***y***_1_, and ***y***_2_ and a distributional assumption about ***x***_*j*_, we approximate the covariance of one variable (***x***_*j*_) with the product of the other two (***w***_2_ = ***y***_1_***y***_2_) as well as the product’s mean and variance.

Repeating this process for each predictor ***x***_*j*_ and following the linear regression equations presented in Section 2.2 allows for calculation of covariate adjusted slope coefficients for the multiple regression model 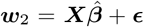 as well as the standard errors of these slope estimates.

#### 2.3.2 Covariance Estimation with the Product of 3 or More Phenotypes

Regression models for larger products of phenotypes can also be approximated by applying the established method recursively: first estimating the covariance of ***x***_*j*_ and ***w***_2_, then leveraging the covariance of ***x***_*j*_ and ***w***_2_ and ***x***_*j*_ and ***y***_3_ to estimate the covariance of ***x***_*j*_ and ***w***_3_, and so forth. This recursion procedure is described in more detail in the appendix and software to carry it out is discussed in Section 2.8.

### 2.4 Binary Phenotypes

While nothing in the previous sections precludes the use of the method on the product of binary phenotypes, some improvements to the method can be made in these cases.

#### 2.4.1 Changes to Estimations

The covariance of two binary phenotypes is estimated using the same general framework as developed in Section 2.3.1. The only changes are to the variance estimates. Instead of estimating a phenotype’s conditional variance from a linear model’s residual variance, we estimate it as

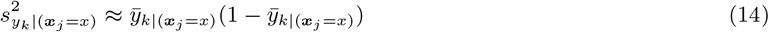

Further, we calculate the product’s sample variance as

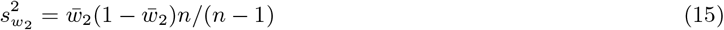

#### 2.4.2 Products as Logical Combinations

Binary phenotypes are of particular importance because their products can be interpreted as logical combinations.

We can represent the logical conjunction ***y***_1_ ∧ ***y***_2_ (read as “***y***_1_ and ***y***_2_”) as the product ***y***_1_***y***_2_. Likewise, we express the logical disjunction ***y***_1_ ∨ ***y***_2_ (“***y***_1_ or ***y***_2_”) as **1**_*n*_ − ((**1**_*n*_ − ***y***_1_)(**1**_*n*_ − ***y***_2_)).

By framing both disjunctions and conjunctions in terms of phenotype multiplication, we can apply our established methods to approximate the covariances of these combinations with predictors and ultimately estimate linear models for these logical combinations.

While the case of the conjunction is a trivial application of the above methods of multiplying phenotypes, we will briefly describe how to model the disjunction. To do so, we consider the modified phenotypes 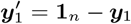 and 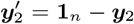. (These represent the statements “not ***y***_1_” and “not ***y***_2_.”) This gives us 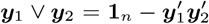. Then, 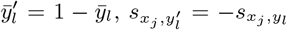, and 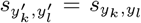. If we set 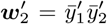, our method allow us to estimate 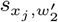 for each ***x***_*j*_ as well as 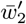 and 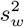. Leveraging these estimates, 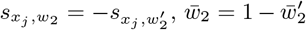, and 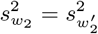, where ***w***_2_ is equivalent to the disjunction ***y***_1_ ∨ ***y***_2_. Using these terms as inputs for the framework presented in Section 2.2 allow for coefficient and standard error estimation for the linear model 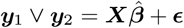.

### 2.5 Simulation

#### 2.5.1 Simulation 1: Type I Error Maintenance

To verify that our linear model with PCSS approach appropriately maintained the Type I error rate at a variety of *α* thresholds, we carried out a simulation under the null hypothesis that the predictor variant has no linear association with any of the phenotypes of interest. This null hypothesis represents a reasonable subset of the exact null hypothesis which is that the *product* of phenotypes has no linear relationship with the predictor. We carried out this simulation with varying sample size, MAF, phenotype means, phenotype correlations, and for continuous phenotypes, phenotype variances, for products of two binary phenotypes, two continuous phenotypes, and three continuous phenotypes with 10^8^ simulations for each collection of continuous phenotypes and 10^7^ simulations for the case of binary phenotypes. Simulation parameters were generated from distributions (details are available in the Appendix in Table S1).

#### 2.5.2 Simulation 2: Comparisons to IPD Models

To evaluate our method’s ability to replicate the results of covariate adjusted linear models fit to IPD, we carried out three 2^*k*^ factorial simulations—one for the product of two binary phenotypes, one for the product of two positive continuous phenotypes, and one for the product of three positive continuous phenotypes. We carried out 1000 simulations at each possible combination of parameters. In each simulation, we modeled the phenotype product as a function of a SNP and binary covariate. For the simulations with only two phenotypes, we also included a continuous covariate in our models.

In all simulations, we simulated *n* subjects’ SNP minor allele counts ***x***_1_ at HWE with varying MAF. We simulated a binary covariate ***x***_2_ with log odds of success *α*_2_***x***_1_. When generating sets of two phenotypes we also generated a continuous covariate ***x***_3_ from a linear regression model with ***x***_1_ with correlation *α*_3_, then centered and standardized. This resulted in a SNP with two covariates (*p* = 3) in our two phenotype simulations, and a SNP with one covariate (*p* = 2) in our three phenotype simulation.

We generated individual phenotype measures through the model

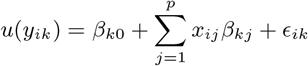

where *u*(*y*_*ik*_) = *y*_*ik*_ for continuous phenotypes, *u*(*y*_*ik*_) = logit(*y*_*ik*_) for binary phenotypes, and 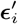 follows a multivariate normal distribution with ***µ*** = **0** and **Σ**_(*i,j*)_ = *σ*_*i*_*σ*_*j*_*ρ*_*ij*_. In all simulations, parameter values were selected such that, under optimal settings, empirical power was roughly 80–90% at a significance threshold of 10^−8^. Full details of simulation parameters are available in the Appendix in Table S2.

In each simulation, we found coefficients, standard errors and two-sided *p*-values for the null hypothesis that there was no relationship between the product of phenotypes and the SNP (***x***_1_) after adjusting for covariates. Values were computed using IPD and PCSS.

Additionally, when simulating two binary phenotypes we fit covariate-adjusted logistic regression models for the logged odds that *y*_1*i*_*y*_2*i*_ = 1 using IPD and returned the relevant two-sided *p*-value to compare the results of the linear model fit using PCSS to the correctly specified logistic model.

### 2.6 Real Data Application

#### 2.6.1 Fatty-Acid Conversion Ratios

Fatty acids are of broad importance for a wide range of cardiometabolic traits (Imamura et al., 2020), with ratios of fatty acids often used as a proxy for conversion efficiency. Previous genome wide association studies have explored the genetic architecture of fatty acids and their ratios (Kalsbeek et al., 2018; Lemaitre et al., 2011; N. L. Tintle et al., 2015; N. Tintle et al., 2020). We modeled 12 fatty acid ratios using both IPD and PCSS using data from the Framingham Heart Study’s Generation-3 and Offspring cohorts downloaded from dbGaP (Mailman et al., 2007).

The 12 ratios can be found in the first column of Table 3. Appendix Table S3 lists all fatty acids used in at least one of the 12 ratios alongside their abbreviations.

Quality control measures included setting Mendelian inconsistencies as missing and excluding SNPs with HWE *p* < 0.00001, MAF < 0.05, or missing values for over 10% of subjects. We excluded individuals missing over 10% of their genetic data after initial quality control and then took a subset of unrelated participants. After quality control we were left with 362,330 SNPs over 1455 individuals (657 from the Offspring cohort and 888 from the Generation-3 cohort).

In addition to the standard PCSS described in Section 2.1, we assumed access to pre-computed means and variances of the reciprocal of each fatty acid as well as the correlation between any fatty acid reciprocal and any other fatty acid, covariate, or SNP to model these ratios using PCSS.

We analyzed each fatty acid ratio through the linear model: Ratio ∼ SNP + age + sex for each SNP in our sample using both IPD and PCSS and tested each SNP for statistical significance with the Bonferroni adjusted threshold *α* = 1.37 × 10^−7^.

### 2.7 Statistical Analysis

#### 2.7.1 Simulation Analysis

To analyze the results of our Type I Error simulations we calculated the empirical Type I Error rate when approximating linear models using PCSS at each specified significance threshold.

For all three 2^*k*^ factorial simulations, we assessed our PCSS method’s errors relative to models fit using IPD when estimating slope coefficients, standard errors, and *t* statistics as well as the test-decision disagreement rate between the IPD and PCSS approaches at a variety of significance thresholds.

We modeled errors in slope coefficients, standard errors, and test statistics through multiple regression models with logical indicators for each of the *k* parameter settings as predictors, testing at the Bonferroni adjusted significance threshold of 0.05/*k*. We also calculated the overall mean bias and variance.

We compared test decisions regarding the significance of the SNP when modeling the phenotype product and adjusting covariates. Test decisions were computed at significance thresholds 10^−1^, 10^−2^, …, 10^−8^. When analyzing binary phenotypes we also compared test decisions between the linear model fit using PCSS and the logistic regression model fit on IPD to demonstrate the robustness of linear models to model binary outcomes. We reported test disagreement rates between the tests using PCSS and IPD at each significance threshold.

#### 2.7.2 Real Data Analysis

We measured our overall bias in slope, standard error, and test statistic estimates as well as the variance of each of these errors for each fatty acid ratio evaluated. We recorded test decisions for both the IPD and PCSS models and recorded which SNPs were found to have significant associations with a given fatty acid ratio. When one approach found a SNP to be significant and the other did not, we recorded if the non-significant result was “borderline” significant (*α* ≤ *p* < 10*α*).

### 2.8 Software

Software to perform these model approximations as well as those developed in Wolf et al. (2020) is available through the R package pcsstools, available on GitHub at https://github.com/jackmwolf/pcsstools.

## 3 Results

### 3.1 Simulation 1

Empirical Type I error rates when using PCSS are displayed in Table 1. In all simulations, the approach’s empirical Type I error rate was below the tested significance threshold.

**Table 1:**
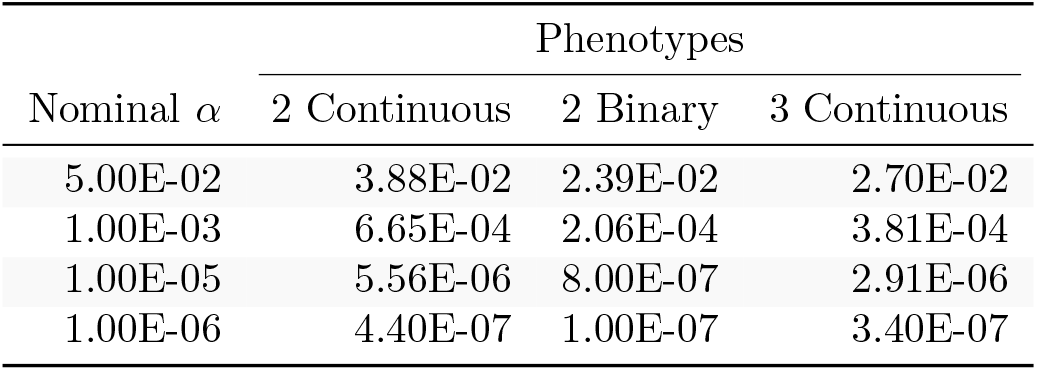
Simulation studies of Type I Error Estimates when testing the linear association between a single SNP and a product of phenotypes using pre-computed summary statistics at significance thresholds: *α* = 0.05, 0.001, 10^−5^, and 10^−6^. Each entry represents the proportion of *p*-values smaller than *α* when modeling the linear relation between a SNP and a product of phenotypes while adjusting for covariates using summary statistics.

### 3.2 Simulation 2

The PCSS method’s errors when approximating slope coefficients, their standard errors, and test statistics are available in Table 2. When aggregated over all simulation settings we observe (small, but) anti-conservative bias in its slope and test statistic estimates in each simulation. The magnitude of the mean test statistic error is comparable across all three simulations. Figure 2 displays our PCSS method’s approximated slope coefficients compared to slope coefficients calculated using IPD for the SNP while modeling the phenotype product and adjusting for covariates. Similar graphical comparisons of standard error and test statistic estimates are available in the Appendix in Figures S2 and S3.

**Table 2:**
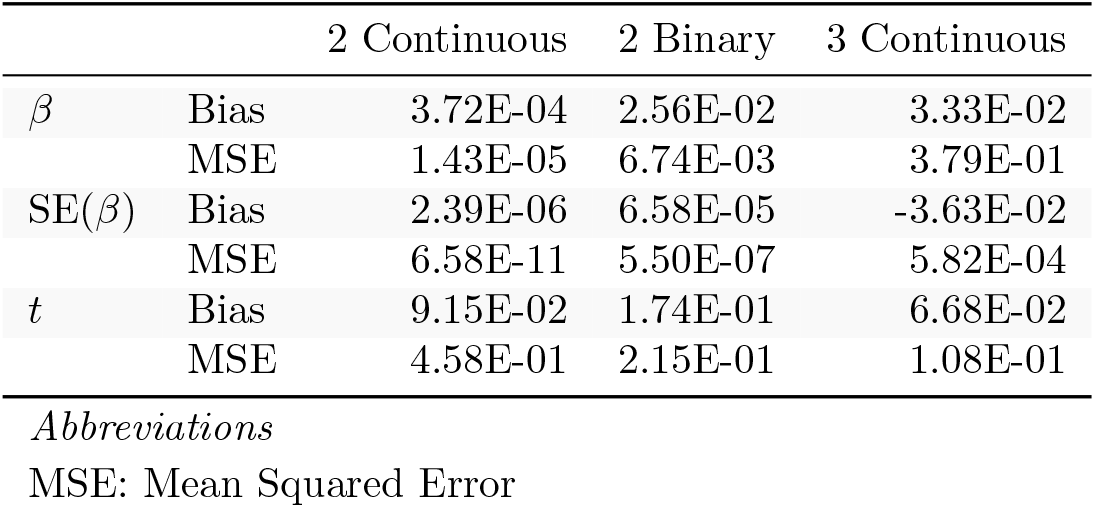
Simulation study approximating a linear model for a product of phenotypes using summary statistics. Summaries of errors when approximating slopes, slope standard errors, and *t*-statistics for a SNP while adjusting for covariates compared to values obtained when calculating these statistics using subject-level data. Errors are calculated as the value calculated using pre-computed summary statistics minus the value found using individual participant data.

**Figure 2:**
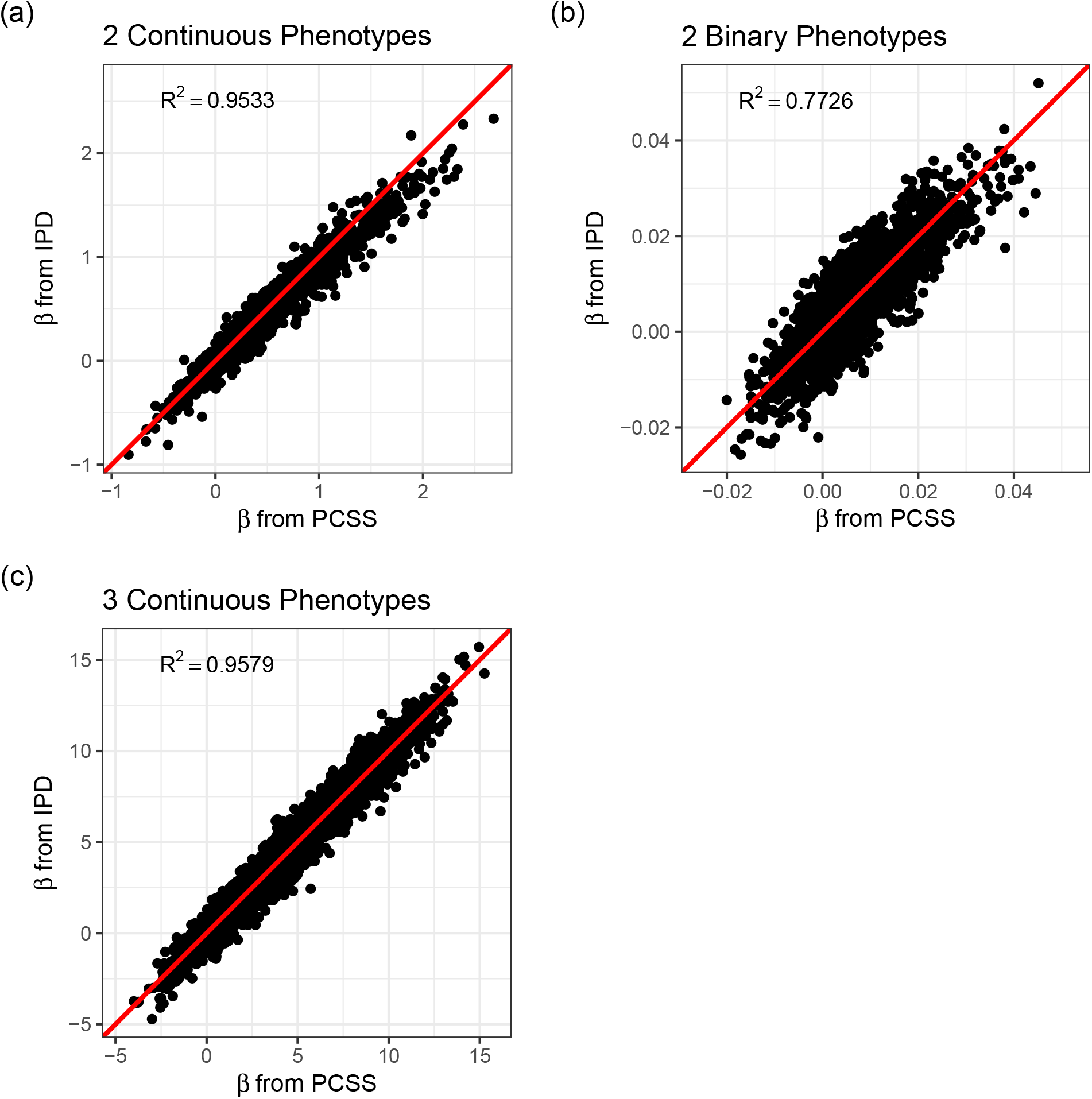
Simulation study approximating a covariate adjusted linear model for a product of phenotypes using pre-computed summary statistics (PCSS) and individual participant data (IPD). Approximated slope coefficients for the SNP while adjusting for covariates are compared to their values when computed using subject-level data.

When modeling estimation errors for two continuous phenotypes through a linear regression model with indicator variables for all of the simulation settings (*k* = 12, *n* = 2^*k*^ × 10^3^), our model for the slope error found all settings except the residual phenotype variances, 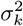, to be significantly associated with the PCSS model’s slope estimate’s error at the adjusted significance threshold 0.05/*k*. All settings had significant associations with our error when estimating the standard error of the slope coefficient, or the test statistic. In the case of two binary phenotypes (*k* = 14, *n* = 2^*k*^ × 10^3^), we found all settings to have significant associations with the error in slope, standard error, and test statistic estimates. For three continuous phenotypes (*k* = 13, *n* = 2^*k*^ × 10^3^), we also found all settings to have significant associations with the error when predicting the slope coefficient, its standard error, and its test statistic.

Figure 3 shows comparisons of estimated and calculated *p*-values for a two-sided *t* test under the null hypothesis that the SNP had no linear association with the phenotype product after adjusting for covariates. Figure 4 shows various error rates rate between the IPD and PCSS models’ test decisions based on these *p*-values at differing significance thresholds. We see that all PCSS models overall disagreement rates to their IPD companions decrease as the significance threshold becomes more stringent. Likewise, when the IPD model rejected the null hypothesis, the PCSS model rarely failed to reject with error rates at most 13% which again decreased as the significance threshold decreased. When the IPD model failed to reject the null hypothesis, the PCSS approaches’ conditional error rates varied by the model’s response. When modeling the product of two continuous or binary phenotypes, the error rate stayed relatively constant across all thresholds at around 3% and 15%, respectively. But, when modeling the product of three continuous phenotypes, the error rate increased as the significance threshold became more strict. Lastly, we can see that when compared to the test decisions of a covariate adjusted logistic regression model, our PCSS approximation of the related linear model tends to reach the same conclusions, with a moderate conservative tendency, especially at more strict significance thresholds.

**Figure 3:**
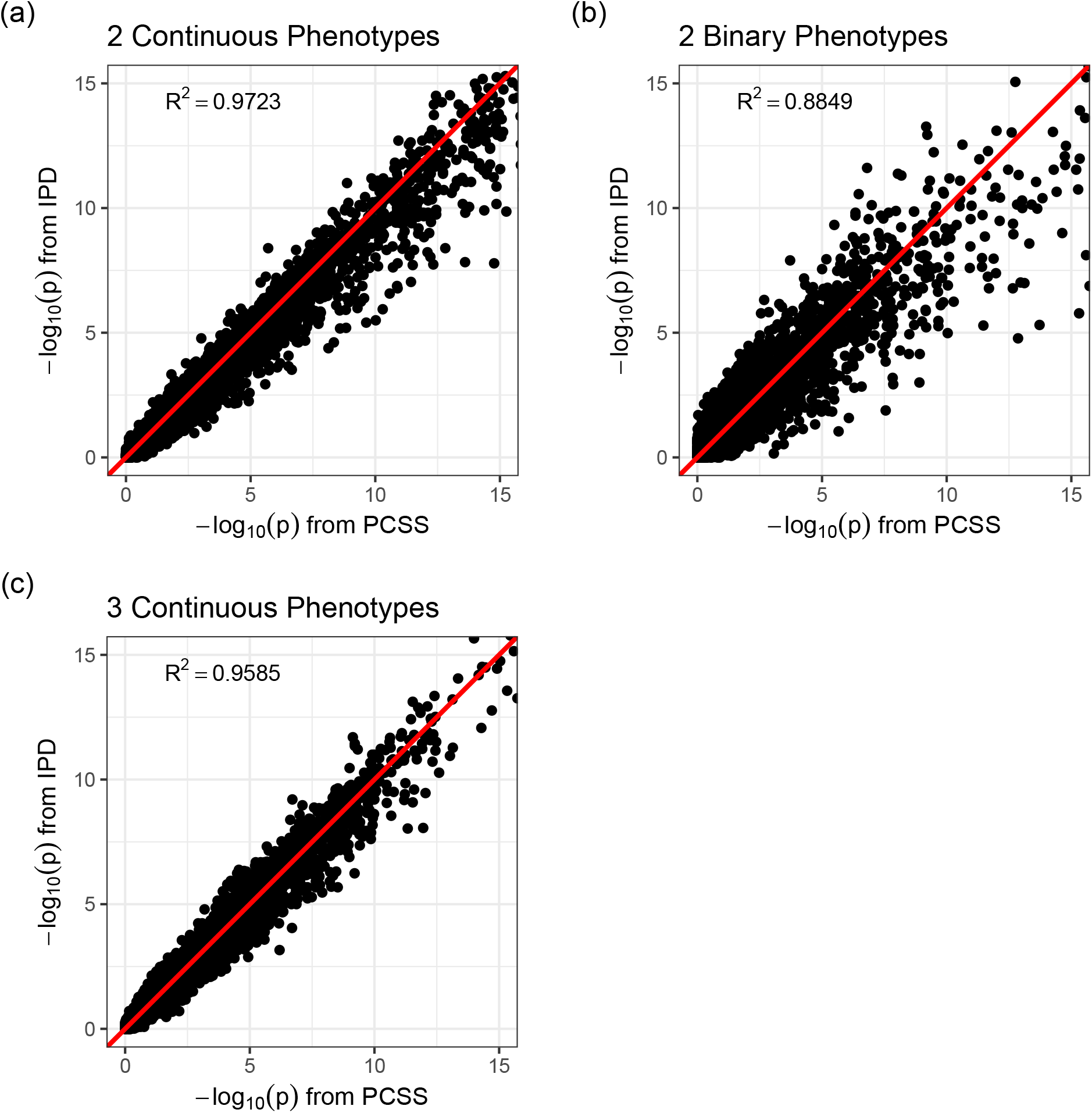
Simulation study approximating a covariate adjusted linear model for a product of phenotypes using pre-computed summary statistics (PCSS) and individual participant data (IPD). Two-sided *p*-values were computed for the null hypothesis that the SNP had no linear effect on the phenotype product while adjusting for covariates.

**Figure 4:**
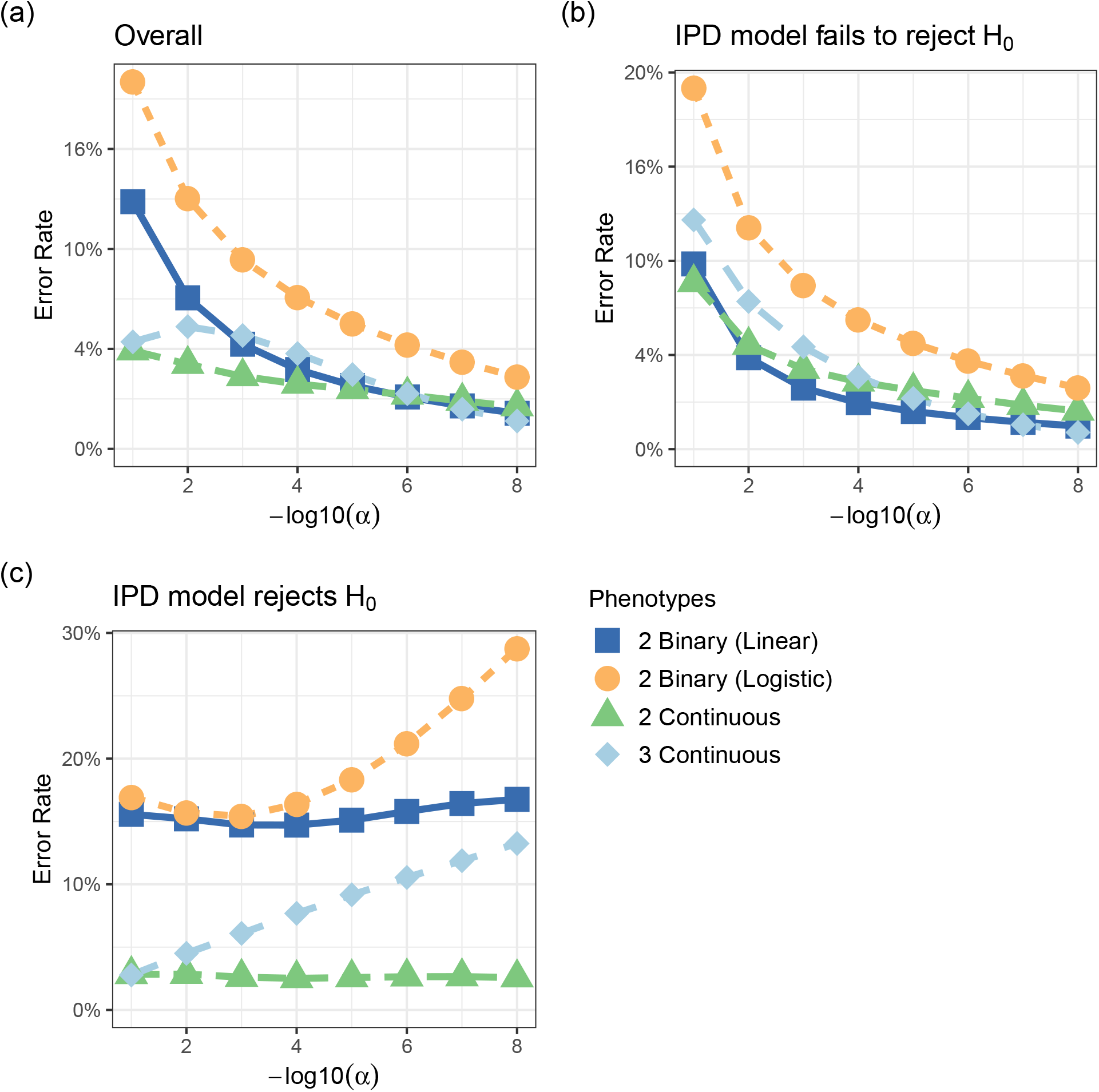
Simulation studies’ test decision error rates evaluating the significance of a SNP in a linear model for a product of phenotypes while adjusting for covariates using Individualized participant Data (IPD) and Pre-Computed Summary Statistics (PCSS) at various significance thresholds (*α*). Comparisons were also made between a logistic regression model fit using IPD on the product of two binary phenotypes and the PCSS model approximating the linear relationship. (a) Percentage of times the IPD and PCSS models’ test decisions disagreed. (b) Error rate conditional on the IPD model failing to reject the null hypothesis. (c) Error rate conditional on the IPD model rejecting the null hypothesis.

### 3.3 Real Data Application

Across all fatty acid ratio models we again observed anti-conservative bias in our slope and test statistic estimates. Our mean slope error was −2.93 × 10^−3^ (Mean Squared Error 0.114) while the mean slope estimate when using IPD was −1.3 × 10^−3^. Our mean test statistic error was −3.34 × 10^−4^ (4.34 × 10^−2^). Values are broken down by ratio in Table 3.

**Table 3:** Summary of errors when approximating the linear model: FA Ratio ∼ snp + age + sex using individual participant data (IPD) and pre-computed summary statistics (PCSS). Each fatty acid ratio was modeled across 362,330 SNPs from 1455 subjects in the Framing Heart Study’s Offspring and Generation-3 cohorts. Errors are calculated as the approximation using PCSS − the value obtained using IPD.

| Ratio | $\beta$ | | | SE($\beta$) | | $t$ | |
| --- | --- | --- | --- | --- | --- | --- | --- |
|  | Mean<sup>†</sup> | Bias | MSE | Bias | MSE | Bias | MSE |
| PA:POA | 5.58E-02 | 1.49E-02 | 4.18E-02 | -4.80E-02 | 2.08E-04 | 1.15E-02 | 1.81E-02 |
| PA:SA | 5.83E-05 | -2.13E-05 | 3.09E-07 | 1.43E-04 | 1.87E-09 | -3.72E-03 | 8.72E-03 |
| POA:OA | -2.08E-05 | -2.10E-06 | 8.08E-09 | -2.69E-05 | 4.69E-11 | -5.83E-03 | 1.90E-02 |
| SA:OA | -8.15E-05 | -1.19E-05 | 3.62E-07 | -3.33E-05 | 3.36E-10 | -1.61E-03 | 8.40E-03 |
| LA:GLA | -7.35E-02 | 2.03E-02 | 1.32E+00 | -5.10E-01 | 1.95E-02 | 1.90E-03 | 3.48E-02 |
| LA:DGLA | 2.04E-03 | 1.99E-04 | 3.58E-04 | -1.84E-03 | 3.12E-07 | 7.86E-04 | 4.54E-02 |
| GLA:DGLA | 4.58E-05 | 1.04E-05 | 2.64E-07 | -7.62E-06 | 1.13E-10 | 4.35E-03 | 7.37E-02 |
| DGLA:AA | -1.99E-05 | -1.54E-06 | 2.84E-08 | 3.79E-05 | 1.09E-10 | -4.31E-04 | 1.38E-02 |
| AA:DTA | 1.76E-03 | -4.20E-04 | 7.21E-05 | -3.76E-03 | 9.42E-07 | -6.40E-03 | 2.74E-02 |
| EPA:DPA_N3 | 2.29E-04 | -1.38E-05 | 1.31E-06 | -4.04E-04 | 1.02E-08 | -1.18E-03 | 5.41E-02 |
| DTA:DPA_N6 | -1.49E-03 | 1.66E-04 | 7.46E-04 | -1.16E-02 | 8.24E-06 | -1.24E-03 | 1.75E-01 |
| DPA_N3:DHA | -4.12E-04 | 6.03E-05 | 3.45E-06 | -1.52E-04 | 2.53E-09 | 5.95E-03 | 4.22E-02 |
| Overall | -1.30E-03 | 2.93E-03 | 1.14E-01 | -4.80E-02 | 2.12E-02 | 3.34E-04 | 4.34E-02 |

*Abbreviations*
MSE: Mean Squared Error
<sup>†</sup> Mean of point estimates when using individual participant data

**Table 4:** Summary of test decisions for a real data application calculating the linear model Fatty Acid Ratio ∼ snp + age + sex using Individual Participant Data (IPD) and Pre-Computed Summary Statistics (PCSS) across 362,330 SNPs from subjects in the Framing Heart Study’s Offspring and Generation-3 cohorts. Cell values are counts conditional on the event listed in the topmost row. Significance threshold of *α* = 1.37×10^−7^.

| Ratio | IPD Significant |  |  | PCSS Significant |  |  |
| --- | --- | --- | --- | --- | --- | --- |
|  | Total | PCSS Sig. | PCSS BL | Total | IPD Not Sig. | IPD BL |
| PA:POA | 0 | 0 | 0 | 0 | 0 | 0 |
| PA:SA | 0 | 0 | 0 | 0 | 0 | 0 |
| POA:OA | 0 | 0 | 0 | 0 | 0 | 0 |
| SA:OA | 6 | 6 | 0 | 9 | 3 | 3 |
| LA:GLA | 5 | 2 | 2 | 2 | 0 | 0 |
| LA:DGLA | 9 | 9 | 0 | 10 | 1 | 1 |
| GLA:DGLA | 8 | 8 | 0 | 8 | 0 | 0 |
| DGLA:AA | 18 | 18 | 0 | 19 | 1 | 1 |
| AA:DTA | 0 | 0 | 0 | 0 | 0 | 0 |
| EPA:DPA_N3 | 0 | 0 | 0 | 1 | 1 | 1 |
| DTA:DPA_N6 | 5 | 4 | 1 | 4 | 0 | 0 |
| DPA_N3:DHA | 11 | 11 | 0 | 11 | 0 | 0 |
| Overall | 62 | 58 | 3 | 64 | 6 | 6 |

*Abbreviations*
BL: Borderline Significant: ( $\alpha \leq p < 10\alpha$ )

Table 3.3 summarizes the number of SNPs found significant when modeling using both IPD and PCSS across all 12 × 362,330 models. We see that of the 93% (58/62) of the time when an IPD model found a SNP to have a significant association with a given fatty acid ratio, the PCSS model also found the SNP to be significant. Moreover, 98% (61/62) of the time when the IPD model found a significant SNP, the PCSS model found the same SNP to have a *p*-value less than 10*α*. Conversely, of the 64 occasions when the PCSS model found a SNP to have a significant association with a given fatty acid ratio, only 6 (9%) occurred when the IPD model did not find the SNP to be significant. On all of these occasions, the IPD model’s *p*-value was less than 10*α*.

## 4 Discussion

We have developed a method that approximates the covariance of products of phenotypes with other variables using only bivariate and univariate pre-computed summary statistics (PCSS). We then demonstrated how this covariance estimation can be used to approximate linear models for products of phenotypes, how these can model logical “and” and “or” statements and how these models can include researchers choice of covariates. We demonstrated our approximation method’s accuracy relative to models fit on individual participant data through multiple simulations and applications to real genetic data.

The approximations shown here show good performance overall. There is a slight tendency towards anti-conservatism, however the Type I error is maintained. Areas of caution in application of the method include potential compounding of errors when applied to products of *m* phenotypes (where *m* is large), multiplying binary phenotypes that exhibit high negative correlation and when phenotypes take on negative values. Additional simulation studies and methodological improvements are needed in these cases and caution should be exhibited when applying our method in these cases. We also note that our method makes assumptions about the fit of the linear model to the data. While these assumptions are the same as in the corresponding analysis of IPD data (e.g., true underlying linear relationship between ***y*** and ***x***), these assumptions may be more acutely important in our PCSS method.

Application of our method to real data from the Framingham Heart Study showed good performance. In general, we have tried to formulate this PCSS method to only rely on commonly available or easily estimated PCSS. However, in our application we assumed that we had the PCSS for ratios of fatty acids. This may not always be the case in practice, but may suggest that these PCSS may be important to pre-compute to assist downstream analyses of ratios.

A variety of limitations of our work are worth noting. First, we used linear regression for a binary response. Previous applications of PCSS have take this approach (Canela-Xandri et al., 2018), and it is generally robust; however, this approach is less precise than when the underlying relationship is truly linear. While some foundations for a logistic modelling approach were recently proposed by Wu et al. (2021), further work is needed to develop a comprehensive model for logistic regression using PCSS. Second, while our simulation study was comprehensive and we demonstrated our method on real data we note that further testing on simulated and real data is encouraged to explore special cases not considered here (e.g., linear combinations of products, adjusting for clustered/family data, etc.)

The use of PCSS provides numerous advantages over IPD data including computational efficiency and reduced concerns about data privacy. However, substantially improved and flexible methods are needed in order to fully leverage PCSS in customized downstream analyses. Our method allows researchers further customization of analyzed phenotypes by opening the door to multiplicative combinations of phenotypes, including logical combinations of binary phenotypes. Approximations used are reasonable, with near perfect maintenance of the Type I error rate and power in most situations. Further work is needed to apply the method to additional datasets and to expand the method to larger classes of combined phenotypes.

## Acknowledgements

The authors of this work were supported by NIH Grant 2R15HG006915-03 and Dordt University. They would like to thank Martha Barnard, Xueting Xia, Nathan Ryder, and Jason Vander Woude for their help with preliminary stages of this project.

## Data Availability Statement

The simulated data that support the findings of this study are available from the corresponding author upon reasonable request. The real data analyses presented in the current publication are based on the use of study data downloaded from the dbGaP web site under dbGaP accessions phs000007.v29.p10 and phs000342.v20.p13.

## Appendix

### Recursive Covariance Estimation

Let ***w***_*l*_ = ***y***_1_***y***_2_ · ***y***_*l*_ = ***w***_*l*−1_***y***_*l*_. In order to estimate 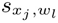 through our established method, we use 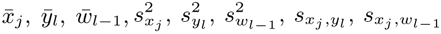, and 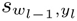 as inputs to the method described in Section 2.3.1. That is, replacing ***y***_1_ with ***w***_*l*−1_ and ***y***_2_ with ***y***_*l*_. While 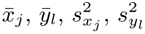, and 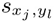 are assumed to be known, we must estimate 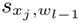 and 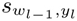.

Continuation of the recursive process starting at *l* − 1 and working down to 2 will yield an estimate for 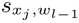, or eventually the base case of 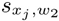.

To approximate 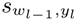, we re-express the term as 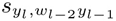. Then, treating ***y***_*l*_ as the predictor (i.e. as we treat ***x***_*j*_), we approximate this term through the method described in Section 2.3.1.

A diagram of the start of this recursion is displayed in Figure S1.

This recursive estimation is impacted by the order in which the phenotypes are multiplied. So, any set of more than two phenotypes will render *m*!/2 possible ways to estimate the regression model through this method (with even more possible through different ways of recursion). Hence, we approximate the covariances and means using all permutations of length *m* of ***y***_1_, …, ***y***_*m*_ unique up to the order of the first two terms as the order of our phenotypes, and take the median of each estimate as the predicted value.

**Figure S1:**
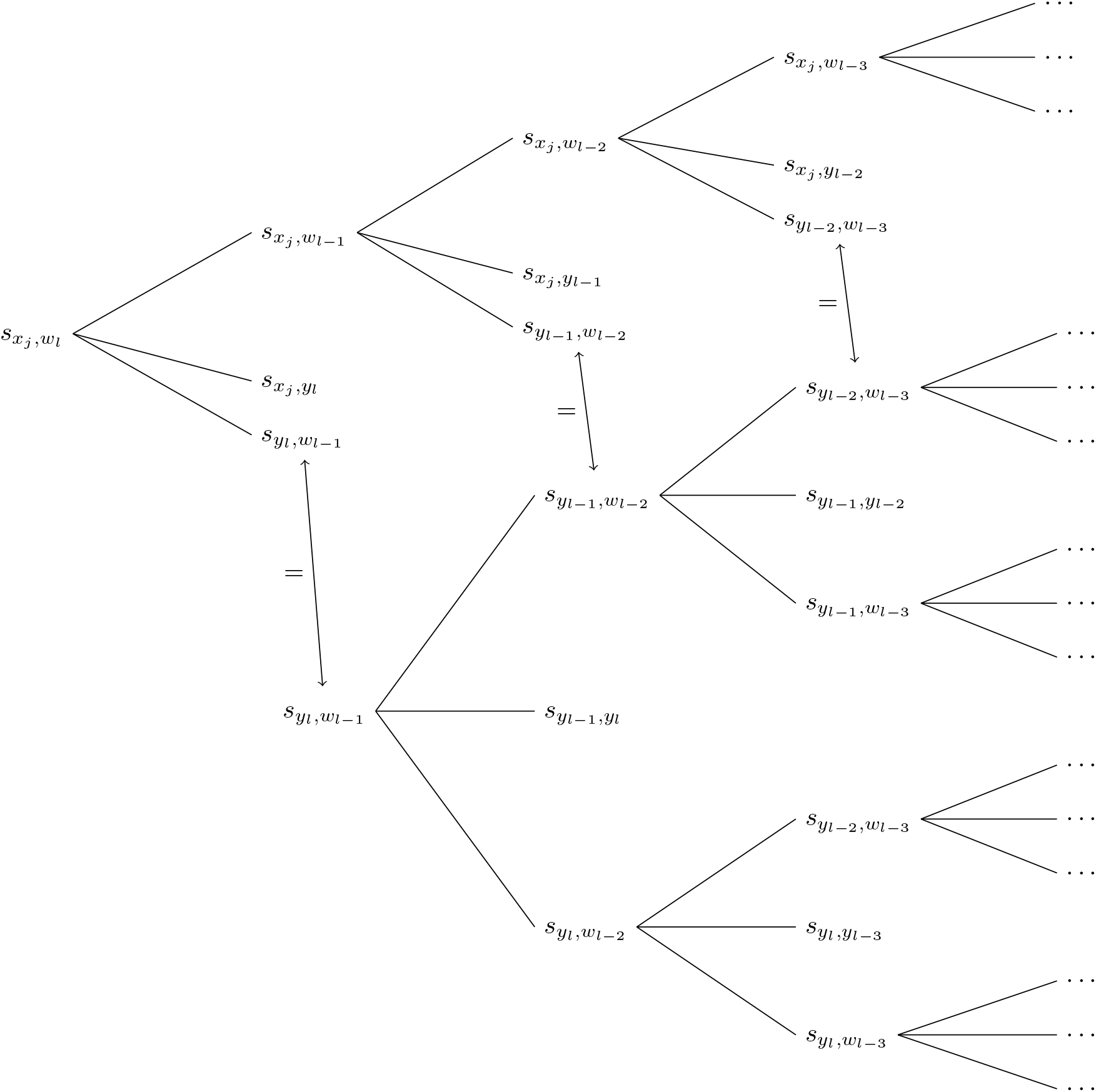
Diagram of the recursive algorithms used to approximate 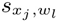. Three covariances are input (along with related means and variances) to approximate their parent node (to the left) using the method established in Section 2.3.1.

**Table S1:** Distributions used to generate simulation parameters for the Type I Error simulations. Continuous phenotypes were generated through a multivariate normal distribution while binary phenotypes were generated through a correlated binomial distribution. Correlations of two binary variables were simulated uniformly from the range of possible correlations for a given set of marginal probabilities *µ*_1_ and *µ*_2_ within the closed interval [−0.25, 0.95].

| | $n$ | MAF | $\mu_k$ | $\sigma_k$ | $\rho_{kl}$ |
| --- | --- | --- | --- | --- | --- |
| Continuous | $10^4, 10^5$ | Unif(0.05, 0.4) | $N(0, 25)$ | Gamma(1, 1/2) | Unif(-0.5, 0.5) |
| Binary | | | Unif(0.2, 0.8) | NA | Unif($f(\mu_1, \mu_2)$) |

**Table S2:** Simulation parameters for 2^*k*^ factorial simulations. We carried out 1,000 simulations at each possible combination of settings for each set of phenotypes. Phenotype measures, or in the case of binary phenotypes logged odds of success, were simulated from a multivariate normal distribution conditional on variables ***x***_1_, ***x***_2_, and, when we generated only 2 phenotypes, ***x***_3_. Parameters were selected such that the empirical power of models using individual patient-level data was around 90% under optimal settings. Columns with a value for Setting 1 but not Setting 2 indicate that the parameter was fixed at the value of Setting 1 in all simulations. Columns with no value for Setting 1 or Setting 2 indicate that the parameter was not used in the simulation.

| | | $n$ | MAF | $\alpha_2$ | $\alpha_3$ | $\beta_{k0}$ | $\beta_{k1}$ | $\beta_{k2}$ | $\beta_{k3}$ | $\sigma_k^2$ | $\rho_{kl}$ |
| --- | --- | --- | --- | --- | --- | --- | --- | --- | --- | --- | --- |
| 2 Continuous<br>($k = 12$) | Setting 1 | $10^4$ | 0.1 | 0 | 0 | 5 | 0.01 | 0.1 | 0.1 | 2 | 0 |
| | Setting 2 | $10^5$ | 0.25 | $\ln 2$ | 0.4 | | 0.05 | 1 | 1 | 3 | 0.4 |
| 2 Binary<br>($k = 12$) | Setting 1 | $10^4$ | 0.1 | 0 | 0 | 0 | $\ln 1.001$ | 0.01 | 0.01 | 1 | 0 |
| | Setting 2 | $10^5$ | 0.25 | $\ln 2$ | 0.4 | $\ln 2$ | $\ln 0.25$ | $\ln 2$ | $\ln 2$ | 5 | 0.4 |
| 3 Continuous<br>($k = 13$) | Setting 1 | $10^4$ | 0.15 | 0 | | 5 | 0.01 | 0.1 | | 1.5 | 0 |
| | Setting 2 | | | $\ln 2$ | | | 0.1 | 1 | | 3 | 0.4 |

*Abbreviations*
MAF: Minor Allele Frequency

**Table S3:**
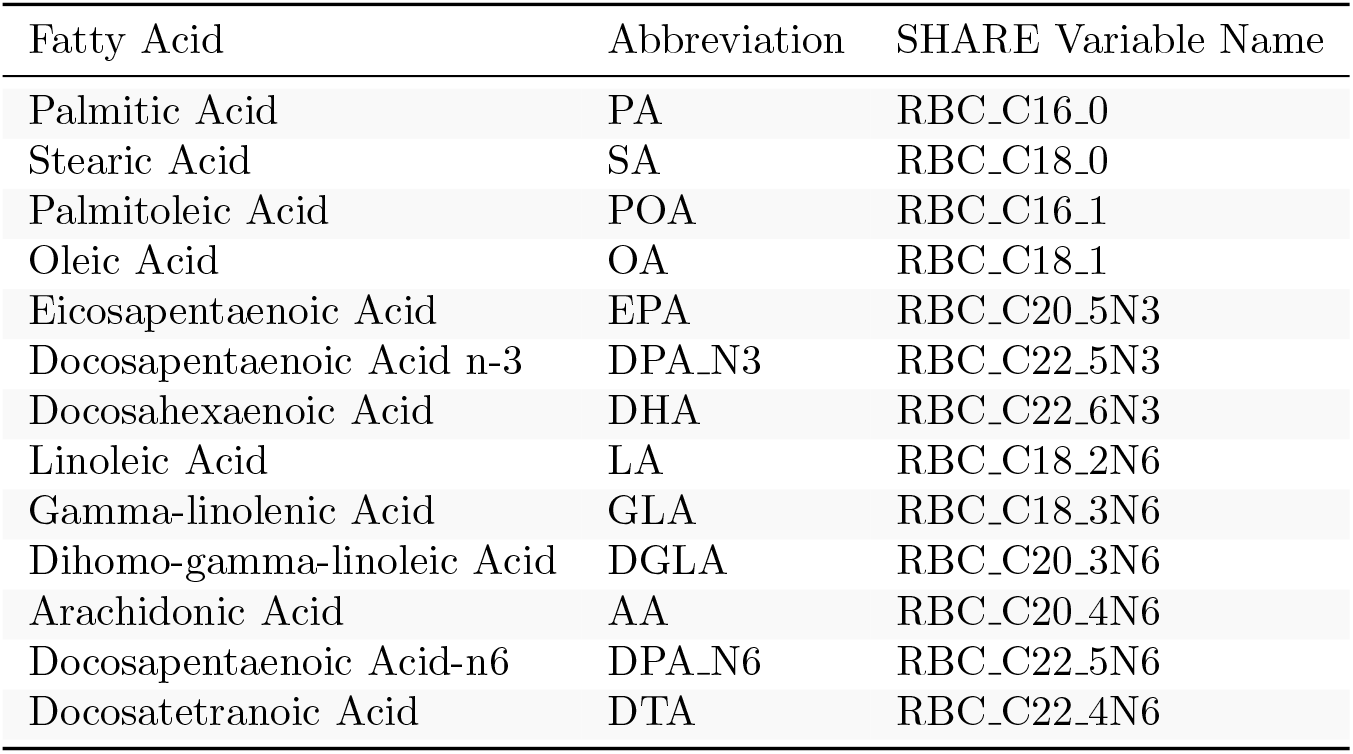
Fatty acids in at least one analyzed ratio with abbreviations.

**Figure S2:**
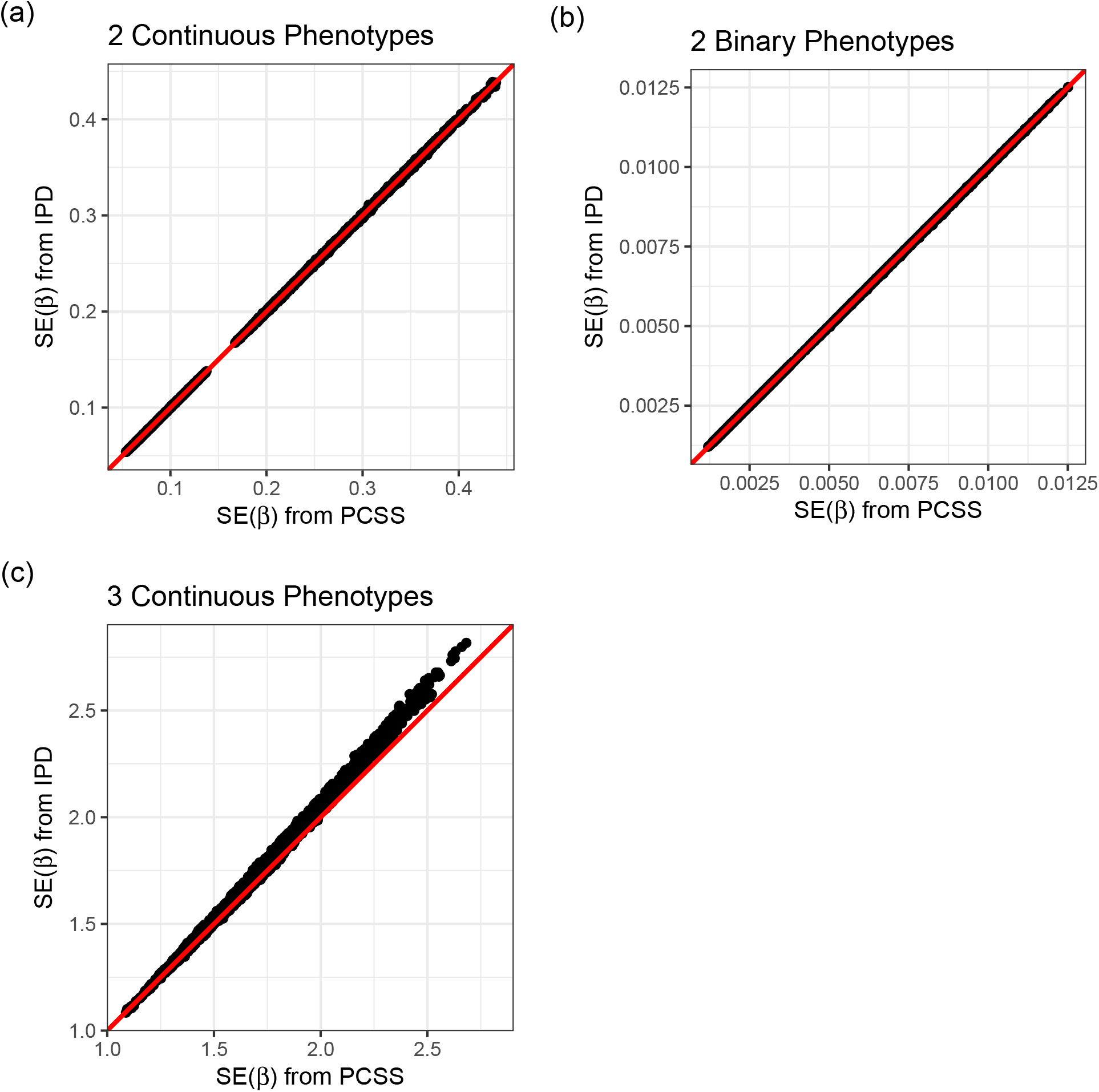
Simulation study approximating a covariate adjusted linear model for a product of phenotypes using pre-computed summary statistics (PCSS) and individual patient data (IPD). Approximated slope standard errors for the SNP while adjusting for covariates are compared to their values when computed using subject-level data.

**Figure S3:**
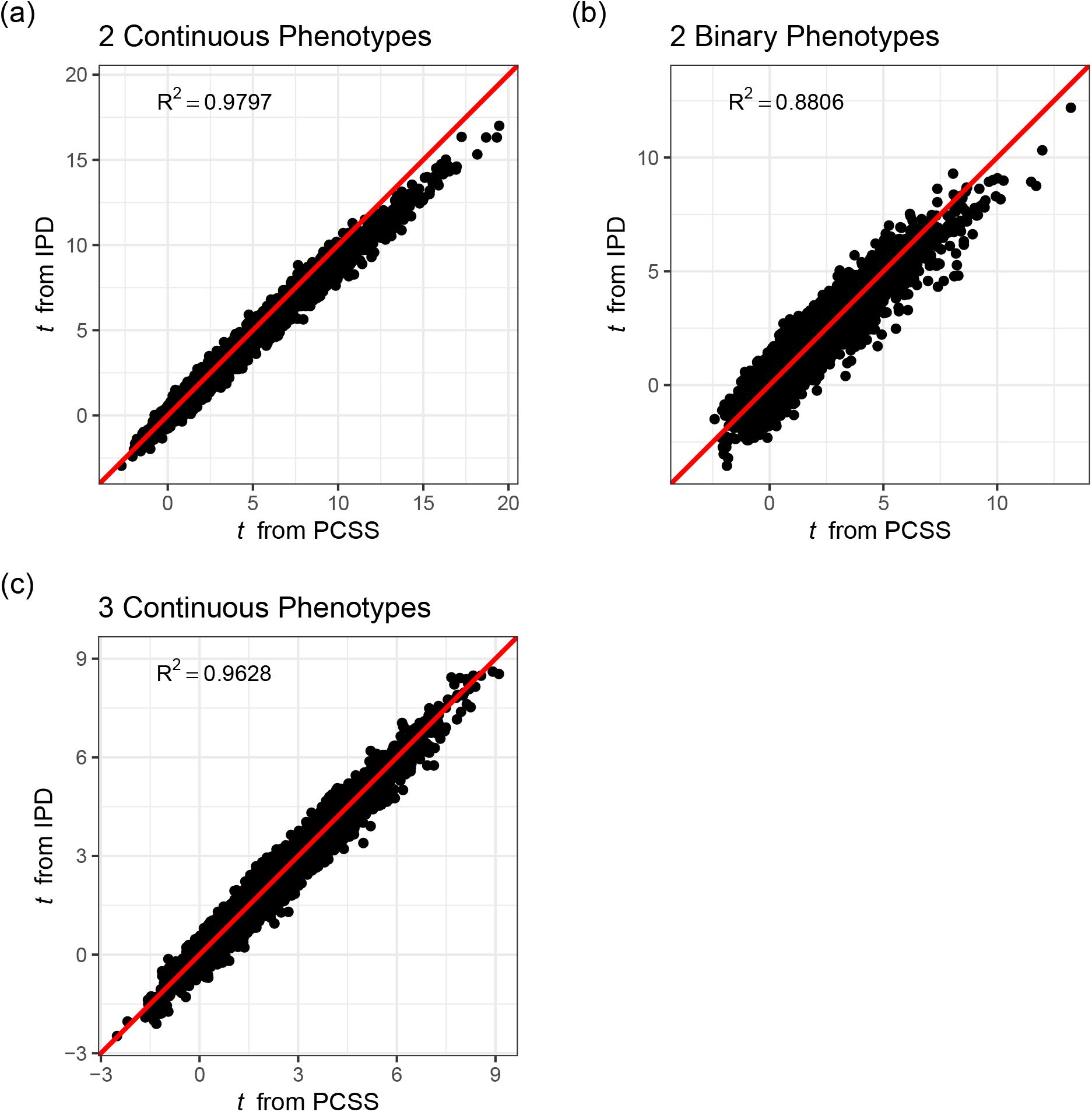
Simulation study approximating a covariate adjusted linear model for a product of phenotypes using pre-computed summary statistics (PCSS) and individual patient data (IPD). Approximated *t* statistics for the SNP while adjusting for covariates are compared to their values when computed using subject-level data.

## References

Baba, K., Shibata, R., & Sibuya, M. (2004, December). Partial correlation and conditional correlation as measures of conditional independence. Australian & New Zealand Journal of Statistics, 46 (4), 657– 664. Retrieved 2020-07-02, from http://doi.wiley.com/10.1111/j.1467-842X.2004.00360.x doi: 10.1111/j.1467-842X.2004.00360.x

Bycroft, C., Freeman, C., Petkova, D., Band, G., Elliott, L. T., Sharp, K., … Marchini, J. (2018, October). The UK Biobank resource with deep phenotyping and genomic data. Nature, 562 (7726), 203–209. Retrieved 2020-12-10, from https://www.nature.com/articles/s41586-018-0579-z (Number: 7726 Publisher: Nature Publishing Group) doi: 10.1038/s41586-018-0579-z

Canela-Xandri, O., Rawlik, K., & Tenesa, A. (2018, November). An atlas of genetic associations in UK Biobank. Nature Genetics, 50 (11), 1593–1599. Retrieved 2020-07-25, from https://www.nature.com/articles/s41588-018-0248-z (Number: 11 Publisher: Nature Publishing Group) doi: 10.1038/s41588-018-0248-z

Cox, N. (2018, October). UK Biobank shares the promise of big data. Nature, 562 (7726), 194–195. doi: 10.1038/d41586-018-06948-3

Diogo, D., Tian, C., Franklin, C. S., Alanne-Kinnunen, M., March, M., Spencer, C. C. A., … Runz, H. (2018, October). Phenome-wide association studies across large population cohorts support drug target validation. Nature Communications, 9 (1), 4285. Retrieved 2020-12-10, from https://www.nature.com/articles/s41467-018-06540-3 (Number: 1 Publisher: Nature Publishing Group) doi: 10.1038/s41467-018-06540-3

Dutta, D., Gagliano Taliun, S. A., Weinstock, J. S., Zawistowski, M., Sidore, C., Fritsche, L. G., … Lee, S. (2019, October). Meta-MultiSKAT: Multiple phenotype meta-analysis for region-based association test. Genetic Epidemiology, 43 (7), 800–814. Retrieved 2020-06-30, from https://onlinelibrary.wiley.com/doi/abs/10.1002/gepi.22248 doi: 10.1002/gepi.22248

Dutta, D., Scott, L., Boehnke, M., & Lee, S. (2019, February). Multi-SKAT: General framework to test for rare-variant association with multiple phenotypes. Genetic Epidemiology, 43 (1), 4–23. Retrieved 2020-06-30, from http://doi.wiley.com/10.1002/gepi.22156 xdoi: 10.1002/gepi.22156

Gagliano Taliun, S. A., VandeHaar, P., Boughton, A. P., Welch, R. P., Taliun, D., Schmidt, E. M., … Abecasis, G. R. (2020, June). Exploring and visualizing large-scale genetic associations by using PheWeb. Nature Genetics, 52 (6), 550–552. Retrieved 2020-07-17, from http://www.nature.com/articles/s41588-020-0622-5 xdoi: 10.1038/s41588-020-0622-5

Gasdaska, A., Friend, D., Chen, R., Westra, J., Zawistowski, M., Lindsey, W., & Tintle, N. (2019). Leveraging summary statistics to make inferences about complex phenotypes in large biobanks. Pacific Symposium on Biocomputing, 24, 391–402. Retrieved 2020-07-10, from https://www.ncbi.nlm.nih.gov/pmc/articles/PMC6417828/ doi: 10.1142/9789813279827_0036

Guo, B., & Wu, B. (2019, July). Integrate multiple traits to detect novel trait–gene association using GWAS summary data with an adaptive test approach. Bioinformatics, 35 (13), 2251–2257. Retrieved 2020-06-30, from https://academic.oup.com/bioinformatics/article/35/13/2251/5201342 xdoi: 10.1093/bioinformatics/bty961

Heatherly, R. (2016, March). Privacy and Security within Biobanking: The Role of Information Technology. The Journal of Law, Medicine & Ethics, 44 (1), 156–160. Retrieved 2020-12-10, from https://doi.org/10.1177/1073110516644206 (Publisher: SAGE Publications Inc) doi: 10.1177/1073110516644206

Imamura, F., Fretts, A. M., Marklund, M., Korat, A. V. A., Yang, W.-S., Lankinen, M., … Forouhi, N. G. (2020, June). Fatty acids in the de novo lipogenesis pathway and incidence of type 2 diabetes: A pooled analysis of prospective cohort studies. PLOS Medicine, 17 (6), e1003102. Retrieved 2020-12-11, from https://journals.plos.org/plosmedicine/article?id=10.1371/journal.pmed.1003102 (Publisher: Public Library of Science) doi: 10.1371/journal.pmed.1003102

Jones, E. M., Sheehan, N. A., Masca, N., Wallace, S. E., Murtagh, M. J., & Burton, P. R. (2012, April). DataSHIELD – shared individual-level analysis without sharing the data: a biostatistical perspective. Norsk Epidemiologi, 21 (2). Retrieved 2020-12-10, from https://www.ntnu.no/ojs/index.php/norepid/article/view/1499 (Number: 2) doi: 10.5324/nje.v21i2.1499

Kalsbeek, A., Veenstra, J., Westra, J., Disselkoen, C., Koch, K., McKenzie, K. A., … Tintle, N. L. (2018, April). A genome-wide association study of red-blood cell fatty acids and ratios incorporating dietary covariates: Framingham Heart Study Offspring Cohort. PLoS ONE, 13 (4). Retrieved 2020-11-17, from https://www.ncbi.nlm.nih.gov/pmc/articles/PMC5898718/ doi: 10.1371/journal.pone.0194882

Kim, J., Bai, Y., & Pan, W. (2015, December). An Adaptive Association Test for Multiple Phenotypes with GWAS Summary Statistics. Genetic Epidemiology, 39 (8), 651–663. Retrieved 2020-06-30, from http://doi.wiley.com/10.1002/gepi.21931 doi: 10.1002/gepi.21931

Lemaitre, R. N., Tanaka, T., Tang, W., Manichaikul, A., Foy, M., Kabagambe, E. K., … Steffen, L. M. (2011, July). Genetic Loci Associated with Plasma Phospholipid n-3 Fatty Acids: A Meta-Analysis of Genome-Wide Association Studies from the CHARGE Consortium. PLOS Genetics, 7 (7), e1002193. Retrieved 2021-01-14, from https://journals.plos.org/plosgenetics/article?id=10.1371/journal.pgen.1002193 (Publisher: Public Library of Science) doi: 10.1371/journal.pgen.1002193

Li, X., Zhang, S., & Sha, Q. (2020, January). Joint analysis of multiple phenotypes using a clustering linear combination method based on hierarchical clustering. Genetic Epidemiology, 44 (1), 67–78. Retrieved 2020-06-30, from https://onlinelibrary.wiley.com/doi/abs/10.1002/gepi.22263 doi: 10.1002/gepi.22263

Mailman, M. D., Feolo, M., Jin, Y., Kimura, M., Tryka, K., Bagoutdinov, R., … Sherry, S. T. (2007, October). The NCBI dbGaP database of genotypes and phenotypes. Nature Genetics, 39 (10), 1181–1186. doi: 10.1038/ng1007-1181

Neale, B. M. (2018). Biobank GWAS. Retrieved 2020-12-10, from http://www.nealelab.is/uk-biobank

Pasaniuc, B., & Price, A. L. (2017, February). Dissecting the genetics of complex traits using summary association statistics. Nature Reviews Genetics, 18 (2), 117–127. Retrieved 2020-06-30, from http://www.nature.com/articles/nrg.2016.142 doi: 10.1038/nrg.2016.142

Ray, D., & Boehnke, M. (2018, March). Methods for meta-analysis of multiple traits using GWAS summary statistics. Genetic Epidemiology, 42 (2), 134–145. Retrieved 2020-06-30, from http://doi.wiley.com/10.1002/gepi.22105 doi: 10.1002/gepi.22105

Simell, B. A., Törnwall, O. M., Hämäläinen, I., Wichmann, H. E., Anton, G., Brennan, P., … Perola, M. (2019, March). Transnational access to large prospective cohorts in Europe: Current trends and unmet needs. New Biotechnology, 49, 98–103. Retrieved 2020-12-10, from http://www.sciencedirect.com/science/article/pii/S1871678418317771 doi: 10.1016/j.nbt.2018.10.001

Tintle, N., Bassett, J., Kuo-Liong, C., Forouhi, N., Kupers, L., Lankinen, M., … Harris, W. S. (2020, March). Circulating Omega-3 Fatty Acid Levels and Total and Cause-specific Mortality: Prospective Evidence From 14 Cohorts in the Fatty Acids and Outcomes Research Consortium. Circulation, 141 (Suppl 1), A43–A43. Retrieved 2021-01-20, from https://www.ahajournals.org/doi/10.1161/circ.141.suppl1.43 (Publisher: American Heart Association) doi: 10.1161/circ.141.suppl_1.43

Tintle, N. L., Pottala, J. V., Lacey, S., Ramachandran, V., Westra, J., Rogers, A., … Shearer, G. C. (2015, March). A genome-wide association study of saturated, mono- and polyunsaturated red blood cell fatty acids in the Framingham Heart Offspring Study. Prostaglandins, Leukotrienes, and Essential Fatty Acids, 94, 65–72. doi: 10.1016/j.plefa.2014.11.007

Wolf, J. M., Barnard, M., Xia, X., Ryder, N., Westra, J., & Tintle, N. (2020). Computationally efficient, exact, covariate-adjusted genetic principal component analysis by leveraging individual marker summary statistics from large biobanks. Pacific Symposium on Biocomputing, 25, 719–730. Retrieved 2020-06-30, from https://www.ncbi.nlm.nih.gov/pmc/articles/PMC6907735/ doi: 10.1142/9789811215636_0063

Wu, P., Wang, B., Lubitz, S. A., Benjamin, E. J., Meigs, J. B., & Dupuis, J. (2021, January). Approximate conditional phenotype analysis based on genome wide association summary statistics. Scientific Reports, 11 (1), 2518. Retrieved 2021-02-10, from http://www.nature.com/articles/s41598-021-82000-1 doi: 10.1038/s41598-021-82000-1

Zhu, X., Feng, T., Tayo, B., Liang, J., Young, J., Franceschini, N., … Redline, S. (2015, January). Metaanalysis of Correlated Traits via Summary Statistics from GWASs with an Application in Hypertension. American Journal of Human Genetics, 96 (1), 21–36. Retrieved 2020-07-05, from https://www.ncbi.nlm.nih.gov/pmc/articles/PMC4289691/ doi: 10.1016/j.ajhg.2014.11.011

